# Working memory modulates auditory perceptual sensitivity during speech planning

**DOI:** 10.1101/2025.06.05.658114

**Authors:** Yaser Merrikhi, Ayoub Daliri

## Abstract

**Purpose:** Working memory plays a critical role in speech production. However, how working memory influences speech systems remains unclear. This study aimed to investigate how maintaining a vowel in working memory alters perceptual sensitivity during speech planning. Method: Thirty healthy adults completed speaking and reading tasks while performing a concurrent working memory task. In speaking blocks, participants produced monosyllabic words, whereas in reading blocks, they silently read them. During each trial, participants categorized an auditory probe stimulus positioned between the vowels /ɛ/ and /æ/. Trials were classified as congruent or incongruent depending on whether the vowel maintained in working memory matched the auditory target. Perceptual sensitivity and speech variability were measured across conditions.

**Results:** In the speaking condition, participants showed higher perceptual sensitivity in congruent trials compared to incongruent trials, indicating that maintaining a vowel in working memory biased auditory target perception during speech planning. No such effect was observed in the reading condition. Additionally, participants exhibited a trend of reduced speech variability in congruent compared to incongruent trials, suggesting that working memory modulates speech outcome. No significant relationship was found between changes in perceptual sensitivity and speech variability across subjects.

**Conclusion:** These findings suggest that maintaining a vowel in working memory shapes auditory target representations and modulates speech outcomes. This study provides novel evidence for the influence of working memory on speech systems.

---

Working memory modulates auditory perceptual sensitivity during speech planning

Working memory, a cognitive system responsible for temporarily storing and manipulating information, is fundamental in enabling goal-directed behavior and communication. It allows individuals to maintain relevant information over short periods and manipulate it to support complex cognitive tasks, including reasoning, learning, and language production (A. Baddeley, 2003). In speech production, working memory enables speakers to plan, structure, and monitor their verbal output dynamically and flexibly (Acheson & MacDonald, 2009; Nozari & Novick, 2017; Schwering & MacDonald, 2020). Deficits in working memory capacity have been widely associated with speech production difficulties, particularly in clinical populations (Archibald, 2017). For example, individuals with Parkinson’s disease often experience disruptions in both working memory and speech, suggesting shared underlying mechanisms affecting speech motor control and cognitive-linguistic processing (L. Altmann & Troche, 2011; Lewis et al., 2003; Manes et al., 2024; McNamara et al., 2008). Empirical studies have further demonstrated that targeted working memory interventions can improve speech production, reinforcing the strong relationship between working memory capacity and speech performance (Delage et al., 2021, 2022). These findings highlight that working memory is not merely a passive storage system but an active and integral component of the speech production process, potentially modulating how speech movements are planned, executed, and monitored.

Theoretical models of working memory provide important insights into how information is stored and manipulated during cognitive tasks. According to Baddeley and Hitch’s influential model (Baddeley & Hitch, 1974), working memory consists of a central executive that supervises two specialized subsystems: the visuospatial sketchpad and the phonological loop. The visuospatial sketchpad temporarily holds visual and spatial information, while the phonological loop specializes in the temporary storage and rehearsal of speech-based information. Research into visuospatial working memory has shown that maintaining visual information can actively modulate the activity of sensory cortices, enhancing neuronal response sensitivity and reducing variability in the visual cortex (Merrikhi et al., 2018a; Merrikhi et al., 2017, 2021). These findings suggest that working memory is not confined to higher-order cortical regions but can directly influence early sensory processing pathways. Additionally, the phonological loop is theorized to maintain speech-like traces in a specialized buffer, which can be refreshed and stabilized through internal rehearsal mechanisms, supporting accurate speech output (Baddeley, 2003). Although computational models of speech production, such as the GODIVA model (Guenther, 2016), have integrated buffering mechanisms to account for speech planning and sequencing, the specific contributions of phonological working memory to sensorimotor control during speech remain poorly understood. There is a critical need to determine how the temporary maintenance of speech representations in working memory interacts with the preparation and execution of speech movements. Addressing this gap is essential for refining theoretical models of speech production and developing targeted clinical interventions for speech and language disorders that involve deficits in working memory.

Theories of speech production emphasize the critical role of two distinct yet interdependent control systems: feedforward and feedback mechanisms (Daliri, 2021; Guenther, 2016; Hickok, 2012; Houde & Nagarajan, 2011; Merrikhi et al., 2025; Parrell et al., 2019; Simonyan, 2014). The feedforward system generates and optimizes motor commands to produce speech movements aimed at achieving specific auditory targets. In parallel, the feedback system continuously monitors the auditory consequences of those movements. Both systems depend on detecting mismatches between the auditory targets and the actual feedback—known as error signals—to make adjustments that maintain the precision of speech production. Therefore, the accuracy of auditory targets would directly impact the generation of error signals and the speech responses generated by the feedforward and feedback control systems. Given that working memory is responsible for maintaining speech targets, it has the potential to shape the representation of auditory targets and contribute to the speech systems. Understanding the contribution of working memory to the feedforward and feedback systems is therefore vital for a comprehensive account of speech motor control.

Inspired by evidence that working memory reduces neural variability in sensory systems (Mitchell et al., 2009; Ruff & Cohen, 2014; Merrikhi et al., 2017, 2018a) and by theoretical models proposing that working memory refreshes and stabilizes speech-related memory traces (Baddeley, 2003), we hypothesize that working memory enhances the precision of auditory targets during speech planning by reducing their representational variability. This increased precision is predicted to improve perceptual sensitivity by making the internal auditory target more distinct and less susceptible to noise or overlap with similar categories. In this framework, when a vowel is about to be produced, it is actively maintained in working memory, its internal representation becomes narrower and more stable, providing a sharper reference for evaluating incoming auditory input. As a result, an ambiguous auditory probe is less likely to be categorized as the planned vowel (i.e., auditory target), since the perceptual system now compares it to a more refined internal standard. Figure 1 illustrates this conceptual model. The black and gray curves represent the internal distributions of auditory targets for the vowels /ε/ and /æ/, respectively. In neutral conditions, internal auditory target distributions are broad and overlapping, leading to greater ambiguity in categorization. Therefore, participants categorize an ambiguous stimulus (such as the boundary stimulus located at the border of two vowels shown by the dashed line in Figure 1) with roughly equal probability as /ε/ or /æ/. However, when the vowel /ε/ is maintained in working memory, the internal distribution for /ε/ becomes narrower (red curve), leading to a lower likelihood of categorizing the boundary stimulus as /ε/ compared to /æ/. This hypothesis positions working memory as an active mechanism that sharpens auditory targets and enhances perceptual discrimination during speech planning.

**Figure 1.**
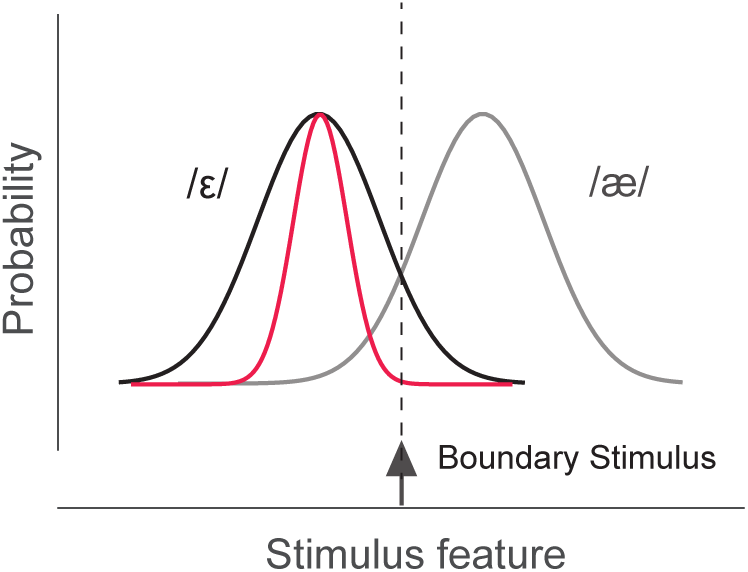
Schematic illustration of how working memory maintenance reduces variability in auditory target distribution. Without working memory engagement, the internal distributions for vowels /ε/ (black) and /æ/ (gray) are broad, leading to similar categorization probabilities for the boundary stimulus (dashed line). When /ε/ is held in working memory, its distribution narrows (red), making the boundary stimulus less likely to be categorized as /ε/.

The primary goal of the present study was to determine the contribution of working memory to speech control systems. As a first step, we investigated how maintaining a vowel in working memory modulates the representation of auditory targets-measured through perceptual sensitivity-during speech planning. To achieve this, we developed a novel experimental paradigm that combined active speech production and reading tasks with a concurrent working memory manipulation. Participants were asked to maintain a vowel in memory while preparing to produce speech or silently read a word, allowing us to compare the influence of working memory under different motor planning demands. By presenting carefully calibrated auditory probe stimuli and measuring perceptual sensitivity, we assessed how working memory maintenance affected participants’ perception of auditory targets. Additionally, we assessed how maintaining a vowel in working memory modulates the variability of speech outcomes. Our approach provides a new framework for examining the dynamic interactions between cognitive and sensorimotor systems during speech production, advancing our understanding of the role of working memory in shaping speech planning and control.

## Method

### Participants

We collected data from 32 healthy adults (21 females, 26 right-handed) with a mean age of 25.5 years (± 2.8 years). We included the participants if they met the following criteria: (1) not having a history of speech, language, psychological, or neurological disorders; (2) being a native speaker of American English; and (3) having a normal hearing within the 250–8000 Hz range (≤ 20 dB HL in all octaves; American Speech-Language-Hearing Association, 1997). The Institutional Review Board at Arizona State University approved the study protocols. All participants provided written informed consent before participating in the study.

### Apparatus

Participants were positioned in front of a computer screen within a sound-isolated booth. A microphone (Shure, SM58) was placed about 15 cm away from the participant’s mouth. The microphone’s output was relayed outside the booth, amplified (TubeOpto 8, ART), converted into digital form, and transferred to a computer through an external audio interface (8pre, MOTU). Using this computer, we displayed the target words on the monitor, and delivered the auditory stimuli to the participants through in-ear headphones (ER-2, Etymotic Research Inc.) following further amplification (S.phone, Samson Technologies Corp). We calibrated the sound levels before each session to maintain the headphone output at 5 dB above the input from the microphone (Merrikhi et al., 2025). For near real-time auditory processing, we utilized Audapter (Cai, 2015), which integrates several signal processing modules for accurate formant analysis and adjustment. Audapter employs linear predictive coding (LPC) for formant identification and uses dynamic programming to enhance tracking accuracy. The specific settings for Audapter included: LPC orders of 15 for female and 17 for male participants, a 48 kHz sampling rate, a downsampling ratio of 3, a 32-sample frame length, and a 5-frame processing lag. We also measured the system’s input-output latency, approximately 18 ms, using a portable digital recorder (Tascam DR-680MKII).

### Procedure

Every participant completed a sequence of tasks, consisting of a practice task, a pre-test task, main tasks (speaking or reading), and sentence reading tasks. The total duration of the experimental session was less than two hours.

### Practice Task

The practice task was designed to familiarize participants with the experimental environment and included 30 trials. During each trial, participants were asked to produce monosyllabic words containing the /ɛ/ vowel (i.e., “hep”, “head”, and “heck”). Each trial lasted 2.5 seconds, with inter-trial intervals lasting 1 to 1.5 seconds. The sequence of target words was shuffled for each trial. Before starting, participants were instructed to prolong the vowel in each word and to keep their vocal intensity steady. We also offered visual feedback to help participants adjust the duration and loudness of the words to the desired parameters (400–600 ms; 70–80 dB SPL).

### Pre-Test Task

This task was designed to assess each participant’s vowel configuration and included 60 trials where participants produced 20 repetitions of monosyllabic target words containing /ε/ and /æ/ within the /hVp/ context. During this task, we did not provide visual feedback on speech duration and intensity unless the participant’s outputs deviated from the desired ranges. Upon completing this task, we calculated the temporal average of the first (F1) and second (F2) formants for each production using the formants estimated by Audapter. For both vowels (/ε/ and /æ/), we automatically determined and visually inspected the centroid of the vowel distribution in the F1-F2 space. Then we used these centroids to estimate the perceptual boundary between /ε/ and /æ/. Our prior research (S. Chao et al., 2019) demonstrated a strong correlation between the production variability of /ε/ and /æ/ and their perceptual boundary. Consequently, we computed the variability-based boundary (VBB) as an estimate of the perceptual boundary. To measure VBB, as shown in Figure 2, we first averaged F1 and F2 for each vowel, we then projected these formant values onto the ε−æ line for each participant, and then calculated the standard deviations (σ_ε_ and σ_æ_) and the distance (D_ε−æ_) between the vowels. We defined VBB in relation to the center of /ε/ with the equation:

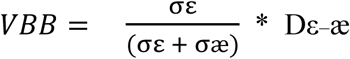

**Figure 2.**
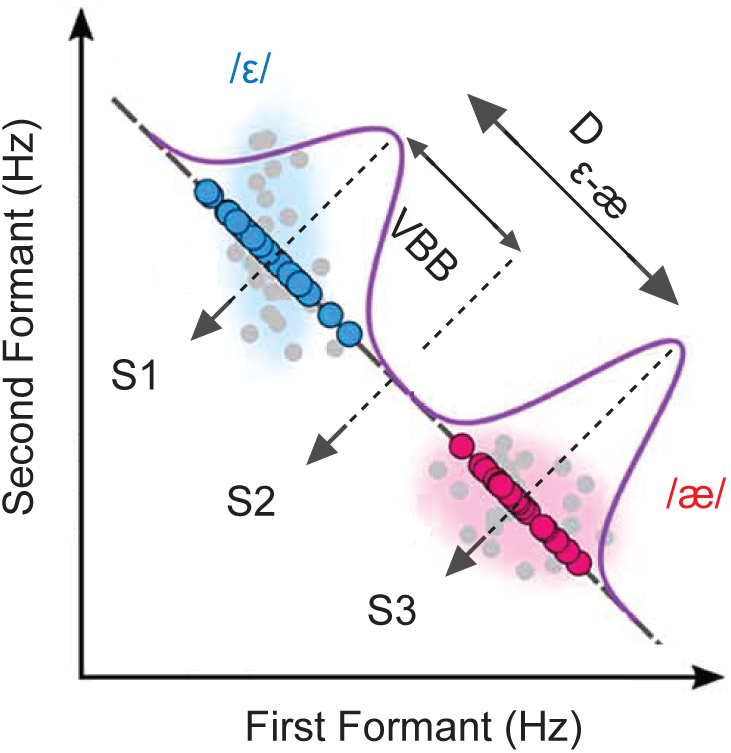
We tailored three probe auditory stimuli required for the main experiments based on each subject’s speech variability (S1: the median of vowel /ε/, S2: located at the boundary between vowels /ε/ and /æ/, S3: the median of vowel /æ/).

Finally, for subsequent experiments, we synthesized three specific auditory probes (Figure 2): S1 centered on the median of vowel /ε/, S2 at the boundary between /ε/ and /æ/, and S3 at the median of vowel /æ/. Each stimulus lasted 50 ms with a 5 ms rise/fall time and was set at 75 dB SPL.

#### Main Task

The main task consisted of two conditions: speaking and reading. We divided the participants into two groups, consisting of 15 and 17 individuals. The first group engaged solely in a speaking condition, completing 8 short blocks of 48 trials each. The second group participated in both speaking and reading conditions, with each condition comprising 6 short blocks of 48 trials. At the start of each block, subjects were briefed about the upcoming condition. As illustrated in Figure 3, during the speaking condition, each trial began by displaying a word in black, which turned green after 600 ms, prompting the subject to produce the word. Conversely, in the reading condition, participants were instructed to silently read the word, eliminating the motor planning aspect of the task (Merrikhi, et al., 2018b). In each trial, one of three probe stimuli (S1, S2, S3) was played to the subjects 400 ms after the target word appeared. These timing details were determined based on our prior research (Daliri & Max, 2015; Merrikhi et al., 2018b). In both the speaking and reading conditions, the words were randomly selected from a collection of six monosyllabic words containing the vowels /ε/ (e.g., “hep”) and /æ/ (e.g., “hap”), with three words representing each vowel. At the end of each trial, regardless of the condition, subjects were required to categorize the probe stimulus by pressing a key corresponding to /ε/ or /æ/.

**Figure 3.**
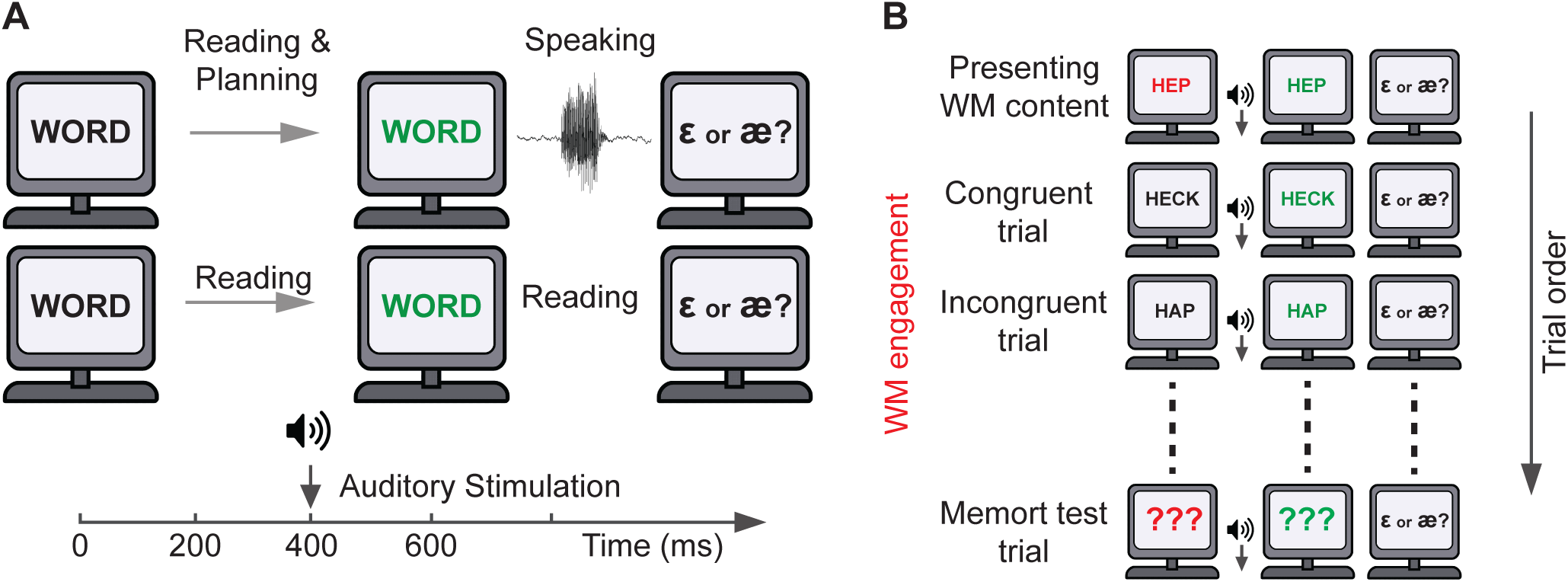
In the speaking condition (A, top row), words initially displayed in black, then turned green after 600 ms, signaling the participant to produce them. In contrast, the reading condition (A, bottom row) required silent reading. In each trial a probe auditory stimulus was played to the subjects 400 ms after the word appeared, and at the end, participants had to categorize the probe by pressing a key corresponding to the vowel /ε/ or /æ/. To engage the working memory (B), the words were displayed in red instead of black. Participants were instructed to memorize the most recent red word, creating congruent and incongruent trials. Finally, participants were asked to verbally reproduce the word they had memorized (memory test trials).

In the first two blocks of each condition, subjects received no working memory instructions (No-WM). In contrast, the subsequent blocks required participants to perform a concurrent working memory task along with their assigned speech production or silent reading tasks (WM blocks). To engage the working memory, in 12% of all trials, we presented the words in red instead of black and instructed the subjects to memorize the most recent red word displayed (Figure 3B). We randomly selected one word at a time from the list of six words as the red word, repeating this process every eight trials until we had selected each of the six words (6 × 8 = 48 trials in each block). Consequently, in 88% of the trials within these blocks, participants continuously held a word in their working memory while engaging in the primary task. This setup yielded two types of trials: congruent and incongruent (Figure 3B). In congruent trials, the vowel maintained in working memory matched the auditory targets in the speaking condition (or corresponded with the word displayed during the reading condition). For simplicity, hereafter, we will also refer to the words displayed in each trial in the reading condition as auditory targets. An exemplar congruent trial might involve memorizing a word with /ε/ while preparing to speak the word /hεp/ (i.e., an auditory target containing /ε/). Conversely, in incongruent trials, the memorized vowel differed from the auditory targets, such as memorizing a word with /ε/ while planning to speak /hæp/. Additionally, during 12% of these trials (termed memory test trials and randomly distributed), participants were asked to verbally reproduce the word they had memorized. Blocks containing errors were repeated to ensure accuracy.

#### Sentence reading Task

To provide participants with a brief break from reading single words and to mitigate experiment-related fatigue, we interspersed a sentence reading task (24 trials) between every two blocks. In each trial, participants read a sentence from the Harvard word list. No visual feedback was given during this task.

### Data analysis

Consistent with our earlier research (Chao et al., 2019; Chao & Daliria, 2023; Daliri, 2021; Daliri & Dittman, 2019), we visually examined formants by superimposing formant trajectories onto the spectrogram of the speech signals. Furthermore, we ensured the correct articulation of target words by replaying all trials. For each subject, we calculated two measures, including perceptual sensitivity and speech variability, which served as the dependent variables in this study. For perceptual sensitivity, we first calculated the percentage of trials in which a participant perceived the S2 probe stimulus (the boundary stimulus) as the planned speech (i.e., auditory target) for each vowel across both congruent and incongruent trials in the speaking and reading conditions, as well as for No-WM trials. We then averaged these response percentages across the two vowels to construct a dependent variable for that participant. For example, in the No-WM trials, we calculated how often a subject perceived the boundary stimulus as the vowel /ε/ in trials where they intended to produce the vowel /ε/ (auditory target). Similarly, we calculated how often they perceived the boundary stimulus as the vowel /æ/ in trials planned for the vowel /æ/ (auditory target). We then averaged these two percentages to determine the percentage of instances in which the subject chose the boundary stimulus as the auditory target during the No-WM trials. Finally, we calculated the normalized perceptual sensitivity of each participant in congruent and incongruent trials by normalizing against their perceptual sensitivity in the No-WM trials, using the following formula:

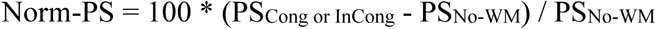

Where Norm-PS is normalized perceptual sensitivity, PS_Cong or InCong_ represent the perceptual sensitivity of participants in congruent or incongruent trials, and PS_No-WM_ is the perceptual sensitivity of participants in the No-WM trials.

To determine the speech variability of subjects under each condition (No-WM, congruent, and incongruent), we first calculated the temporal average of F1 and F2 values within the early time interval (0–100 ms). We then adjusted the speech productions in the F1-F2 space (measured in mel scale) for each condition by subtracting the average speech production of that condition from each production. We then calculated the standard deviation of these adjusted speech productions in the F1-F2 space and used this measure as the speech variability for each respective condition. Finally, we calculated the normalized speech variability of each participant in congruent and incongruent trials by normalizing against their speech variability in the No-WM trials, using the following formula:

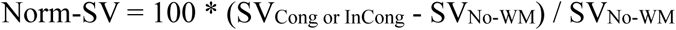

Where Norm-SV is normalized speech variability, SV_Cong or InCong_ represent the speech variability of participants in congruent or incongruent trials, and SV_No-WM_ is the speech variability of participants in the No-WM trials.

All data analyses were performed using MATLAB version R2024b (The MathWorks, Natick, MA, United States), utilizing the Statistics and Machine Learning Toolbox. We conducted paired t-tests to evaluate differences in perceptual sensitivity and speech variability between congruent and incongruent trials within the reading and speaking conditions. We used the Pearson correlation coefficient to study the relationship between changes in perceptual sensitivity and speech variability across subjects.

## Results

### Maintaining a vowel in working memory alters perceptual sensitivity in the speaking condition

Figure 4A illustrates participants’ normalized perceptual sensitivity for congruent (red circles) and incongruent (blue circles) trials in the speaking condition. Error bars indicate the mean ± standard error (SE). In the first group (n = 15), participants were less likely to categorize the boundary stimulus as the auditory target during congruent trials compared to No-WM trials, resulting in negative normalized perceptual sensitivity (Figure 4A, red circles, Norm-PS_Cong_ = -4.66% ± 3.73%). Conversely, in incongruent trials, participants were more likely to perceive the boundary stimulus as the auditory target relative to No-WM trials, yielding a positive normalized perceptual sensitivity (Figure 4A, blue circles, Norm-PS_InCong_ = 4.49% ± 4.63%). Crucially, participants perceived the boundary stimulus significantly less often as the auditory target in congruent trials than in incongruent trials (t(14) = -3.686, *p* = 0.002). In other words, when subjects planned to produce a vowel, they were less likely to perceive the boundary stimulus as that vowel (i.e. auditory target) if they maintained the same vowel in their working memory (i.e. congruent trials) as opposed to when they held a different vowel in their working memory (i.e. incongruent trials).

**Figure 4.**
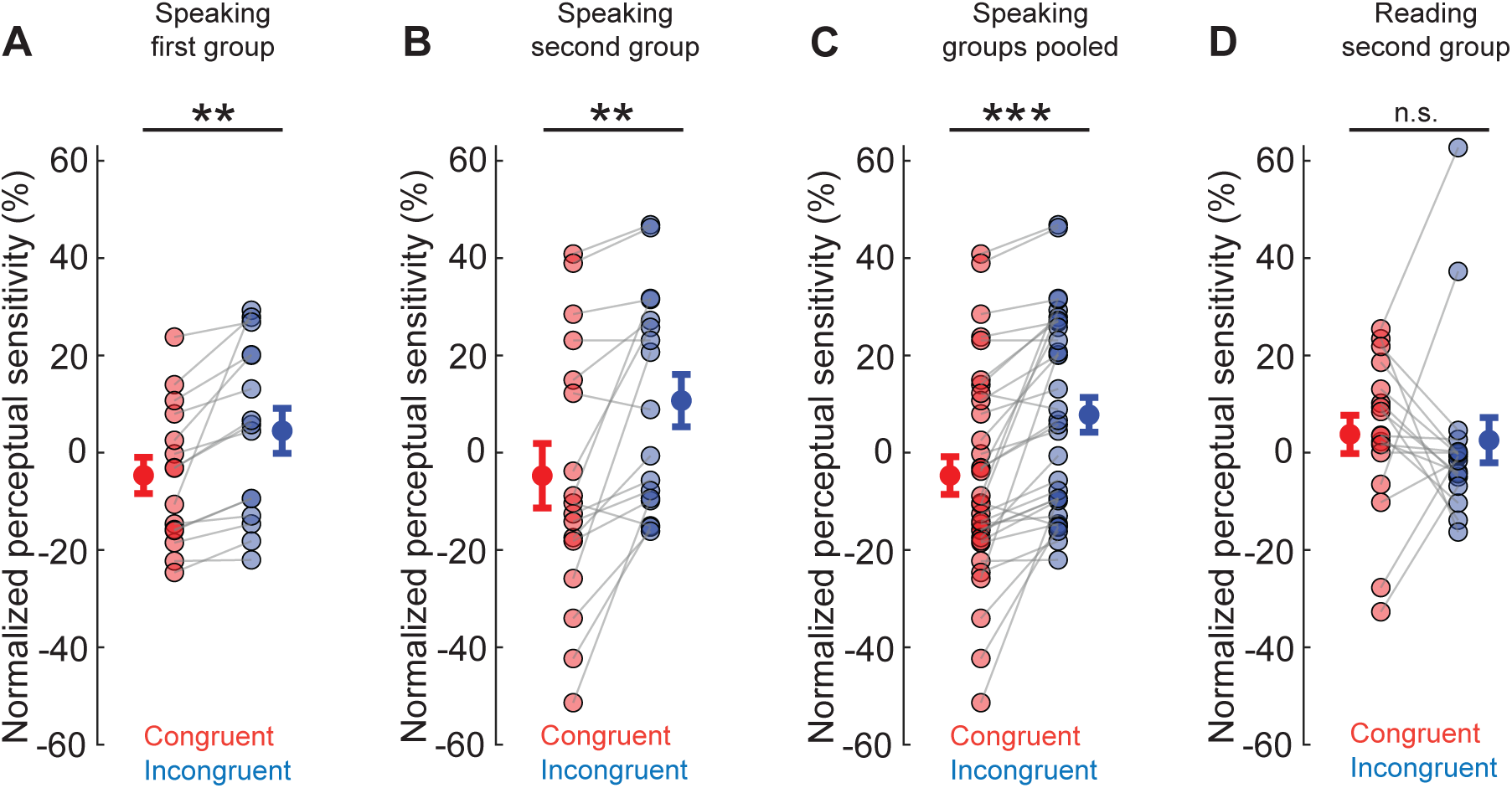
Normalized perceptual sensitivity of participants during speech planning for congruent (red) and incongruent (blue) trials in the speaking condition for the first group (A), second group (B) and both groups combined (C), and in the reading condition for the second group (D).

### Maintaining a vowel in working memory alters perceptual sensitivity in the speaking condition but not the reading condition

Figure 4B presents the normalized perceptual sensitivity of participants in the second group for congruent (red circles) and incongruent (blue circles) trials in the speaking condition. Consistent with the findings from the first group, participants in the second group (*n* = 17) were less likely to perceive the boundary stimulus as the auditory target during congruent trials compared to No-WM trials (Figure 4B, red circles, Norm-PS_Cong_ = -4.73% ± 6.63%). In contrast, they were more likely to perceive the boundary stimulus as the auditory target in incongruent trials relative to No-WM trials (Figure 4B, blue circles, Norm-PS_InCong_ = 10.70% ± 5.36%). Our findings revealed that participants perceived the boundary stimulus significantly less often as the auditory target in congruent trials compared to incongruent trials (t(16) = -3.943, *p* = 0.001).

Figure 4C displays the normalized perceptual sensitivity for all participants combined from the first and second groups (*n* = 32) in the speaking condition, comparing congruent (red circles) and incongruent (blue circles) trials. Participants were less likely to perceive the boundary stimulus as the auditory target during congruent trials compared to No-WM trials (Figure 4C, red circles, Norm-PS_Cong_ = -4.70% ± 3.87%). In contrast, they were more likely to perceive the boundary stimulus as the auditory target in incongruent trials relative to No-WM trials (Figure 4C, blue circles, Norm-PS_InCong_ = 7.79% ± 3.57%). As in the separate group analyses, participants perceived the boundary stimulus significantly less often as the auditory target in congruent trials than in incongruent trials (t(31) = -5.176, *p* < 0.001).

Figure 4D shows the normalized perceptual sensitivity of participants in the second group during the reading condition for congruent (red circles) and incongruent (blue circles) trials. Participants were more likely to perceive the boundary stimulus as the auditory target in both congruent and incongruent trials compared to No-WM trials (Figure 4D, Norm-PS_Cong_ = 3.78% ± 3.94%, Norm-PS_InCong_ = 2.58% ± 4.66%). In contrast to the speaking condition, our findings revealed no significant difference in perceptual sensitivity of participants between congruent and incongruent trials (t(16) = 0.216, *p* = 0.831). In other words, only in the speaking condition, where subjects actively planned to produce a vowel, they perceived the boundary stimulus as an auditory target less frequently in congruent trials, where the planned vowel matched the vowel held in working memory, compared to incongruent trials, where a different vowel was retained in memory. We did not observe this influence of working memory on auditory perception during speech planning in the reading condition, where subjects silently read the word without planning to produce it.

### Maintaining a vowel in working memory influences speech variability

Figure 5 illustrates the normalized speech variability for congruent (red circles) and incongruent (blue circles) trials in the first group (Figure 5A), the second group (Figure 5B), and both groups combined (Figure 5C). For the first group, participants produced slightly less variable speech in congruent trials compared to No-WM trials, resulting in negative normalized speech variability (Figure 5A, red circles, Norm-SV_Cong_ = -0.78% ± 8.76%). In contrast, speech variability increased in incongruent trials relative to No-WM trials, leading to positive normalized speech variability (Figure 5A, blue circles, Norm-SV_InCong_ = 3.65% ± 6.96%). While participants tended to produce less variable speech in congruent than incongruent trials, this difference was not statistically significant (t(14) = -1.203, *p* = 0.248). In the second group, participants exhibited reduced speech variability in both congruent and incongruent trials compared to No-WM trials (Figure 5B, red circles, Norm-SV_Cong_ = -10.91% ± 4.49%; blue circles, Norm-SV_InCong_ = -5.16% ± 4.38%). Our findings revealed that participants produced less variable speech in congruent than incongruent trials, although this difference was not statistically significant (t(16) = -1.81, *p* = 0.087). When pooling data from both groups (n = 32, Figure 5C), participants showed lower speech variability in both congruent and incongruent trials compared to No-WM trials (Figure 5C, red circles, Norm-SV_Cong_ = -6.16% ± 4.75%; blue circles, Norm-SV_InCong_ = -1.03% ± 4.02%). Importantly, participants produced significantly less variable speech in congruent trials than in incongruent trials (t(31) = -2.16, *p* = 0.038).

**Figure 5.**
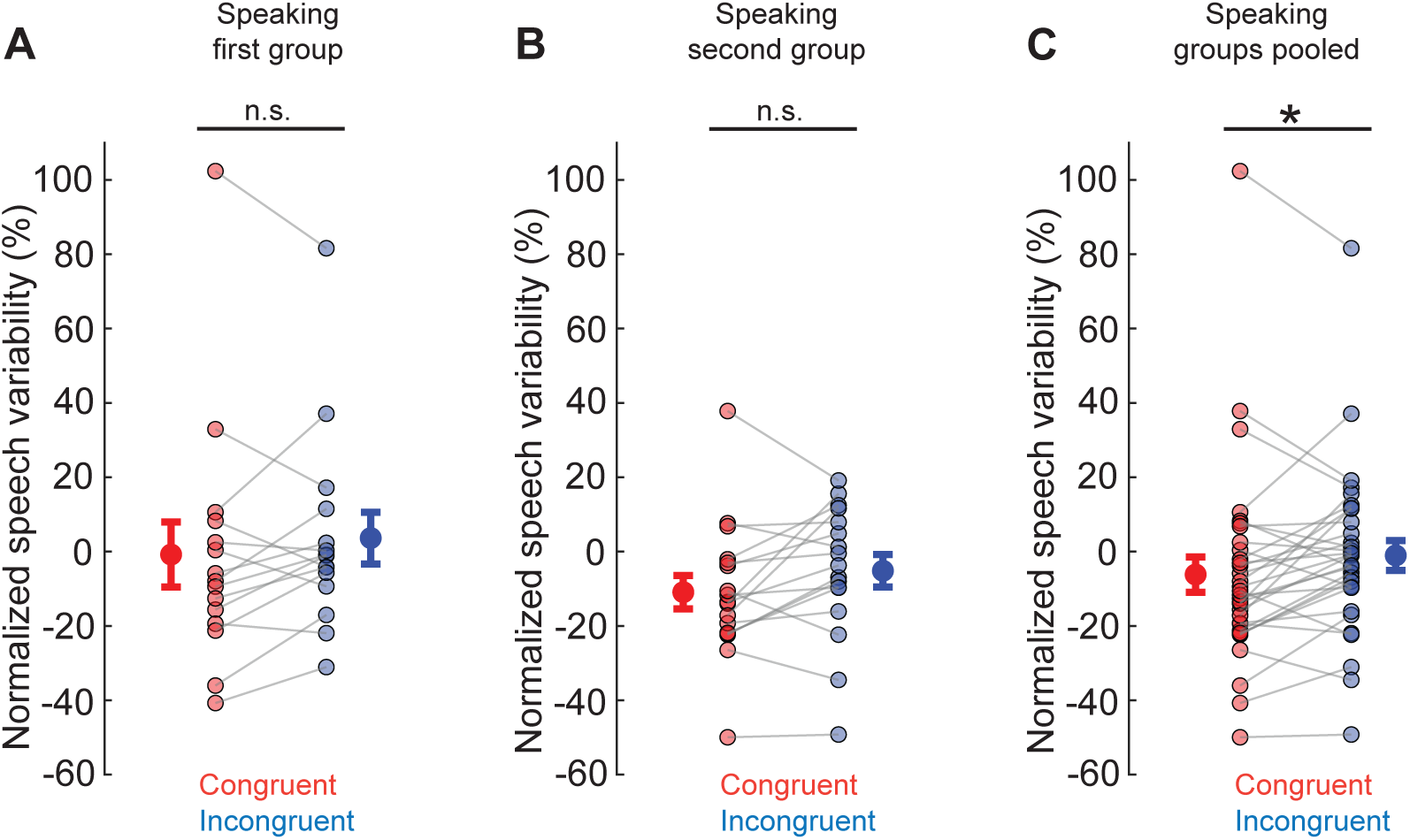
Normalized speech variability for congruent (red circles) and incongruent (blue circles) trials in the first group (A), second group (B) and both groups combined (C).

Finally, we studied whether there is any relationship between working memory-related changes in perceptual sensitivity and speech variability across the subjects (i.e., the difference in perceptual sensitivity or speech variability of subjects between congruent and incongruent trials). We did not find any significant relationship between changes in perceptual sensitivity of subjects and the changes in their speech variability in either first (r = -0.031, *p* = 0.914, Pearson correlation coefficient), second group (r = -0.38, *p* = 0.130, Pearson correlation coefficient) and both group pooled together (r = -0.21, *p* = 0.260, Pearson correlation coefficient).

## Discussion

The primary objective of the present study was to investigate how maintaining a vowel in working memory influences (1) perceptual sensitivity during speech planning and (2) speech motor variability. We developed a paradigm that combined speech production task with a concurrent working memory task. Participants categorized auditory probe stimuli located at the boundary between two vowels while holding either congruent or incongruent vowel information in working memory. We found that (1) working memory modulated participants’ perceptual sensitivity during speech planning but not during reading, and (2) participants exhibited reduced speech variability when maintaining a congruent vowel in working memory compared to an incongruent vowel.

Our findings provide novel evidence that maintaining a vowel in working memory biases perceptual sensitivity during speech planning. Specifically, we found that participants were less likely to categorize the boundary stimulus as the auditory target when the vowel held in working memory matched the vowel they were preparing to produce (congruent trials), compared to when the vowels differed (incongruent trials). In other words, maintaining a specific vowel in working memory appears to enhance the precision (or reduce the variability) of the auditory target representation for that vowel during speech planning. This behavioral finding may reflect a similar mechanism to that observed in neural studies, where working memory content has been shown to reduce the variability of neuronal responses in a spatially-specific manner. For example, neurons exhibit reduced independent and shared variability when the location held in working memory matches the neurons’ receptive fields (i.e., congruent trials), compared to when the remembered location does not match their receptive fields (i.e., incongruent trials) (Cohen & Maunsell, 2009; Merrikhi et al., 2018a; Merrikhi et al., 2017; Mitchell et al., 2009; Ruff & Cohen, 2014). By maintaining a vowel in working memory, participants may gain access to a more reliable (i.e., less variable) internal auditory target representation, making them less likely to categorize an ambiguous external auditory stimulus as the planned vowel. While the present finding is based on behavioral responses in a speech domain and the prior studies examined neural variability in visual working memory, these observations raise the possibility that working memory may serve a broader role in shaping sensory representations across modalities and systems.

A key aspect of our findings is that the influence of working memory on auditory perception was specific to the speaking condition and not observed during reading. Participants did not show differences in perceptual sensitivity between congruent and incongruent trials in the reading condition, where motor planning demands were minimal. This specificity indicates that active motor planning, rather than passive visual processing, is necessary for working memory to shape auditory target representations. These results align with models suggesting that speech planning engages internal sensory predictions and supports the idea that working memory interacts directly with these predictive mechanisms (Guenther, 2016; Houde & Nagarajan, 2011). In the absence of speech motor planning—as in the reading condition—auditory target representations appear more stable and less susceptible to memory-related modulation. Together, these findings suggest that working memory and motor control systems are tightly integrated during goal-directed speech behavior.

In addition to its influence on perception, we found that maintaining a vowel in working memory influenced speech variability. Participants produced speech outcomes with lower variability in congruent trials than incongruent trials when data were pooled across both experimental groups. However, when analyzed separately within each group, this reduction in speech variability did not reach statistical significance. Our task was primarily designed to investigate perceptual sensitivity rather than speech variability, which may explain why the effects on speech output were smaller and more variable. As a result, the task may not have been sufficiently sensitive to detect consistent motor effects at the group level. Moreover, because speech variability was measured within the first 100 ms of vocalization, it was primarily driven by the feedforward control system, with minimal contribution from auditory feedback processes. As such, the current findings provide only a partial view of how working memory might influence speech motor control. Future studies should be specifically designed to examine the impact of working memory on speech variability driven by both speech control systems. Importantly, the modulation of speech motor variability was not correlated with the modulation of auditory target sensitivity, suggesting that working memory may influence perception and production through distinct, though possibly parallel, mechanisms. While maintaining a vowel in working memory appears to sharpen internal auditory target representations (i.e., enhancing perceptual sensitivity), it may independently constrain variability in motor execution. Future studies specifically designed to disentangle these perceptual and motor pathways will be essential to understanding how working memory interacts with different components of the speech production system.

The observed influence of working memory on both perceptual sensitivity and speech variability highlights the dynamic interaction between cognitive and sensorimotor systems during speech production. Theories of speech production propose that feedforward and feedback control systems work together to achieve accurate and fluent speech (Daliri, 2021; Guenther, 2016; Houde & Nagarajan, 2011; Parrell et al., 2019). Feedforward control relies on learned motor programs to generate expected sensory outcomes, while feedback control monitors auditory outputs and corrects deviations from expected sensory outcomes in real-time. Our findings suggest that working memory may contribute to both systems by stabilizing predicted sensory outcomes, thereby promoting more consistent speech output. Maintaining a vowel in working memory may sharpen sensory predictions used by the feedforward system, leading to a more stable speech plan. Simultaneously, enhanced sensory predictions may also improve feedback-based error detection. Although the current study did not reveal a direct correlation between perceptual and motor modulations, these findings support the idea that working memory interacts with multiple components of the speech production system, potentially through distinct but converging mechanisms.

These results also have important implications for theoretical models of speech. Existing models, such as GODIVA (Guenther, 2016), incorporate mechanisms for planning and sequencing speech units but have paid less attention to how working memory maintenance might bias internal sensory representations during speech planning. Our findings suggest that working memory is not a passive buffer but actively shapes sensory predictions and motor outputs, contributing to both the perception and execution of speech. By elucidating how working memory influences speech motor control, our results help refine current models of speech production by integrating cognitive mechanisms—specifically, how memory-driven representations interact with feedforward and feedback processes. More broadly, this research bridges a critical gap in cognitive-linguistic models by revealing how working memory facilitates speech motor planning, thereby deepening our understanding of the cognitive architecture that supports fluent speech.

Our findings may also have clinical relevance. Working memory impairments are prevalent in several clinical populations, including individuals with Parkinson’s disease—groups that often exhibit speech production difficulties (L. Altmann & Troche, 2011; Lewis et al., 2003; Manes et al., 2024; McNamara et al., 2008). Current speech treatments tend to focus on behavioral strategies without accounting for underlying cognitive mechanisms (Allen, 2013; Ingham et al., 2018; Maas et al., 2008; Wren et al., 2018). Our findings suggest that working memory directly shapes auditory target representations and speech variability, supporting the idea that cognitive state influences sensorimotor control. This insight provides a foundation for developing theory-driven, personalized interventions that assess and incorporate an individual’s working memory capacity into treatment planning. By manipulating working memory load during therapy, clinicians may be able to more precisely diagnose cognitive contributions to speech deficits and adapt interventions to each individual’s profile. This personalized approach could lead to more effective and durable improvements in speech intelligibility.

Several limitations of the present study should be acknowledged. First, the task was primarily designed to measure perceptual sensitivity rather than speech variability, which may have limited our ability to detect motor effects, particularly in a relatively small sample. Future research should implement tasks that are specifically optimized to capture subtle variations in speech motor output. Second, we did not include a standardized measure of individual working memory capacity. This decision was partly based on our sample of healthy young adults, who are typically expected to perform within a high and relatively narrow range of such measures. Nevertheless, incorporating working memory assessments in future studies will be important for capturing individual variability and extending the paradigm to broader populations. Third, the current study was limited to healthy young adults, and it remains unclear whether similar effects would generalize to older adults or individuals with speech and language impairments. Fourth, while our behavioral findings offer indirect evidence for interactions between working memory and speech motor systems, future studies employing neuroimaging or electrophysiological methods will be needed to directly examine the underlying neural mechanisms. Finally, this study represents a first step in investigating how working memory influences speech production, and the experimental paradigm will benefit from further refinement and fine-tuning to more effectively isolate the contributions of cognitive and sensorimotor processes.

In sum, this study investigated how maintaining a vowel in working memory during speech planning influences auditory target perception and speech production. We found that working memory modulates auditory target representations during motor planning and stabilizes speech outputs. These results highlight the important role of working memory in shaping sensory and motor processes during speech production and suggest new directions for refining models of speech motor control and working memory integration.

## Acknowledgments

This work was supported by grants from the Arizona Biomedical Research Centre (RFGA2024-022-018) and Arizona State University (ISSR seed grant) awarded to Y. Merrikhi, and grants from the National Institutes of Health awarded to A. Daliri (R01-DC019905 and R01-DC020162). The content is solely the responsibility of the authors and does not necessarily represent the official views of the funding agencies.

## Data Availability Statement

The data set collected for the current study is available upon reasonable request from the corresponding author (Y. Merrikhi).

